# A comparative analysis of 5-azacytidine and zebularine induced DNA demethylation

**DOI:** 10.1101/048884

**Authors:** Patrick T. Griffin, Chad E. Niederhuth, Robert J. Schmitz

## Abstract

The non-methylable cytosine analogs, 5-azacytidine and zebularine, are widely used to inhibit DNA methyltransferase activity and reduce genomic DNA methylation. In this study, whole-genome bisulfite sequencing is used to construct maps of DNA methylation with single base pair resolution in *Arabidopsis thaliana* seedlings treated with each demethylating agent. We find that both inhibitor treatments result in nearly indistinguishable patterns of genome-wide DNA methylation and that 5-azacytidine had a slightly greater demethylating effect across the genome at higher concentrations. Transcriptome analyses revealed a substantial number of up-regulated genes, with an overrepresentation of transposable element genes, in particular CACTA-like elements. This demonstrates that chemical demethylating agents have a disproportionately large effect on loci that are otherwise silenced by DNA methylation.

## Introduction

Cytosine DNA methylation, the covalent addition of a methyl group to the 5’ carbon of a cytosine nucleotide, is required for viability in plants and mammals that possess this base modification. Its presence or absence is known to influence gene expression (Finnegan et al. 1993), heterochromatin status (Mathieu et al. 2007), and genomic integrity through transposon silencing (Saze and Kakutani 2007; Johannes et al. 2009; Reinders et al. 2009). In mammals, DNA methylation covers most of the genome with the exception of certain unmethylated CG dinucleotides “CpG islands” (Kafri et al. 1992), and aberrant DNA methylation is associated with cancer in humans (Ohm et al. 2007; Widschwendter et al. 2007; Gal-Yam et al. 2008). In plants, DNA methylation is distributed differently than in mammals and is found enriched at pericentromeric regions and at lower levels on chromosome ends (Zhang et al. 2006; Zilberman et al. 2007; Cokus et al. 2008; Lister et al. 2008; Niederhuth and Schmitz 2014). Because DNA methylation in flowering plants is meiotically inherited and changes in DNA methylation states can affect morphological variation, it is thought of as a latent reservoir of phenotypic diversity (Ji et al. 2015). Consequently, its manipulation has been pursued in recent years to discover potentially beneficial new traits, particularly in crop species.

The diversity of DNA methylation patterns in plants is attributed in part to the variety of DNA methyltransferase enzymes that establish and maintain it. Methylated CG sites (mCG), regardless of their location in the genome, are faithfully propagated and maintained by DNA METHYLTRANSFERASE 1 (MET1) (Finnegan et al. 1996). CHG methylation (mCHG) is most commonly found in transposons and repeat elements, and it is maintained by a feed forward loop that requires the activity of the DNA methyltransferase CHROMOMETHYLASE 3 (CMT3) and the histone methyltransferase KRYPTONITE (KYP) (Jackson et al. 2002; Cao et al. 2003; Du et al. 2012). Curiously, CMT3 also appears to be involved in the establishment of gene body DNA methylation (gbM), as it was recently discovered that species that have lost CMT3 have no gbM (Bewick et al. 2016). CHH methylation (mCHH) is dependent on either CMT2 or DOMAINS REARRANGED METHYLTRANSFERASE 2 (DRM2). CMT2 largely acts in deep heterochromatin regions of the genomes as well as within the bodies of large transposons (Zemach et al. 2013). In contrast, DRM2, as part of the RNA directed DNA Methylation (RdDM) pathway, methylates mostly at the edges of repeats and transposons in euchromatin (Law and Jacobsen 2010).

The development of whole-genome bisulfite sequencing (WGBS) has advanced the understanding of DNA methylation in plant genomes (Cokus et al. 2008; Lister et al. 2008). A comprehensive analysis of WGBS on *Arabidopsis thaliana* mutants defective for DNA methylation helped describe the specific roles of RdDM-associated enzymes and siRNA-independent DNA methylation enzymes, while also establishing the interplay between the two pathways (Stroud et al. 2013). These and other studies continue to provide a valuable resource to plant researchers interested in the mechanistic underpinnings of how DNA methylation is established and maintained in plant genomes.

The chemical inhibition of DNA methyltransferases has been utilized as a transient alternative to study the effect of DNA methylation loss in plants (Pecinka and Liu 2014). Two of the most widely used chemical demethylating agents, 5-azacytidine (AZA) and zebularine (ZEB), act as non-methylable cytosine analogs; incorporating into the DNA double helix in the place of cytosine with each cycle of DNA replication (Pecinka and Liu 2014). Previous studies have shown that AZA covalently binds to DNA methyltransferases, forming nucleoprotein adducts, which depletes the number of active DNA methyltransferase enzymes in the cell (Jones and Taylor 1980; Creusot et al. 1982; Christman et al. 1983; Santi et al. 1984). ZEB, a more stable alternative to AZA, inhibits DNA methylation in a similar manner, although it is not thought to form an irreversible bond with DNA methyltransferases (Champion et al. 2010). Although AZA and ZEB have been widely used in plants, a genome-wide, comprehensive analysis of either chemical on DNA methylomes has been missing.

In this study, we use WGBS to compare the genome-wide effects of AZA or ZEB treatment on *A. thaliana* seedlings. Although each demethylating agent seems to have an indiscriminate, concentration-dependent effect genome-wide, AZA may be more effective at higher concentrations. mCG was found to be proportionally less impacted by AZA in comparison with mCHH in both the pericentromeres and chromosome arms. RNA-seq was performed to identify potential effects of chemical demethylation on gene expression. Transposable element genes were by far the most highly upregulated class, in particular CACTA-like elements. Genes with high amounts of methylation in all contexts were more highly upregulated than those categorized as gene body-methylated genes. The results of this study help to further clarify the effect of non-methylable cytosine analogs on DNA methylation genome-wide and will provide a guide for future application of these tools.

## Methods

### I. Seed sterilization, plate preparation, and chemical treatments

Agarose gel (Ameresco) with added half-strength Linsmaier and Skoog nutrients (Caisson Laboratories, Inc) was prepared and autoclaved. 5-azacytidine (Sigma) and zebularine (APExBIO) were dissolved in dimethyl-sulfoxide (DMSO) and water, respectively, before being added to the liquefied cooling agar at final concentrations of 25 μM, 50 μM and 100 μM. Columbia-0 (Col-0) background *A. thaliana* seeds were subjected to an ethanol-based seed sterilization and approximately 30 seeds plated per treatment. As a control, seeds were plated on agar containing DMSO with no chemical demethylating agent (AZA mock-treated control), or agar containing neither DMSO nor chemical demethylating agent (untreated control). After a two-day stratification period at 4°C, the seeds were transferred to room temperature and allowed to grow for 8 days under constant light.

### II. DNA extraction and Whole-Genome Bisulfite Sequencing

*A. thaliana* seedlings from each agar plate were pooled and DNA was extracted using the DNeasy Plant Mini Kit (QIAGEN). MethylC-seq libraries were prepared as previously outlined (Urich et al. 2015). Briefly, sonicated DNA (sheared to ≈200-400 bp) was selected with Ser-Mag Speed Beads (Thermo Scientific) and then subjected to end-repair, A-tailing, and adaptor ligation. The DNA was then treated with sodium bisulfite from the EZ DNA Methylation Gold Kit (Zymo Research) and the bisulfite converted DNA was PCR-amplified for 10 cycles. After cleanup of the PCR product, the DNA libraries were sequenced using the Illumina Next-Seq 500 at the Georgia Genomics Facility. One sample from each treatment group and control group was deeply sequenced, with average coverage of 23.0 to 28.1 (**Table S1**). Downstream analysis was carried out on FASTQ files that were mapped to the TAIR10 reference genome after being trimmed for adaptors and preprocessed to remove low quality reads using Methylpy (Schultz et al. 2015b; Schultz et al. 2015a). A second replicate of control samples and seedlings treated with 100 μM of AZA or ZEB were lightly sequenced (**Table S2**) and run through the web tool *FASTmC*, a tool for genome-wide estimation of DNA methylation levels (Bewick et al. 2015).

Genome-wide methylation levels from deeply-sequenced samples were calculated using weighted methylation (Schultz et al. 2012). BedTools (Quinlan and Hall 2010) was used to make windows of consistent sizes across the genome and extract methylation data for genomic features. Custom scripts in Python were used to calculate methylation levels from windows produced by Bedtools. Custom scripts in R were used to rearrange data sets and visually represent the data. Further, the linear model function in R, lm(), was used to determine the association between demethylating agent concentration and DNA methylation levels (e.g. lm(weighted methylation ~ concentration) (**Table S3**).

*A. thaliana* genes were classified as gene-body methylated (gbM), mCHG-enriched, or mCHH-enriched using a previously defined list (Niederhuth et al. 2016). Briefly, genes were tested for enrichment of mCG, mCHG, and mCHH sites in coding sequences against a background methylation rate using a binomial test (Takuno and Gaut 2012). Genes enriched for mCG, but not mCHG or mCHH, were classified as gbM genes. Genes enriched for mCHG, but not mCHH were classified as mCHG-enriched genes. These genes can also contain mCG, which is often found alongside mCHG. mCHH-enriched genes were those genes enriched for mCHH, but could also contain mCG and mCHG (Soppe et al. 2000; Niederhuth et al. 2016).

### III. RNA-seq

Col-0 seeds were treated with 100 μM of 5-azacytidine alongside DMSO mock-treated controls as before. RNA was extracted using TRIzol (Thermo Scientific) and RNA libraries were made using the TruSeq Stranded mRNA Library Prep Kit (Illumina). Three replicates of AZA-treated seedlings and four replicates of mock-treated seedling were then sequenced using the Illumina Next-Seq 500 instrument at the Georgia Genomics Facility. Reads were mapped to the TAIR10 *A. thaliana* reference genome using the default settings of Tophat 2 version 2.0.14 (Kim et al. 2013). Cuffdiff software version 2.2.1 with default settings was used to calculate expression levels and identify differentially expressed genes (Trapnell et al. 2013). To eliminate infinite expression differences, 0.1 was added to every expression value, and the log2-fold-changes between treated and untreated samples were calculated. P-values were corrected for multiple testing using Benjamini-Hochberg False Discovery Rate (q value). Genes were considered differentially expressed with a *q* < 0.05 and a log_2_-fold-change greater than 2.0 or less than −2.0.

In assessing the enrichment of upregulated methylation-categorized (e.g gbM, mCHG genes, etc) and transposable element genes, each subgroup was subjected to a Fischer’s Exact Test via the fisher.test() function in R. Methylation-categorized gene categories were tested as a subset of all genes, whereas transposable element gene categories were tested against all transposable element genes.

## Results

To assess the effect of AZA and ZEB, a sample of each treatment was deeply sequenced using WGBS and the genome-wide methylation level for all cytosines in each context of DNA methylation were plotted (**Figure 1A**, **Table S2**). A concentration-dependent decrease in DNA methylation was observed in both AZA and ZEB-treated DNA. The relationship between DNA methylation and chemical concentration was highly correlated (all *p* values < 0.05) for DNA treated with either AZA (r^2^= 0.99) and ZEB (r^2^= 0.88) for all methylated cytosines, and for CG (AZA r^2^= 0.99; ZEB r^2^= 0.93) and CHG methylation (AZA r^2^= 0.96; ZEB r^2^= 0.99) specifically (**Table S3**). This observation suggests that at 100 μM, neither the effect of AZA nor ZEB on CG or CHG methylation is saturated. CHH methylation did not substantially decrease between the 50 μM and 100 μM treated samples for either AZA or ZEB-treated seedlings, suggesting some amount of saturation. Consequently, the relationship between inhibitor concentration and methylation loss was less highly correlated (AZA r^2^= 0.65; ZEB r^2^= 0.61). A second replicate of seedlings treated with 100 μM AZA and ZEB using low coverage sequencing combined with *FASTmC* analysis confirmed the genome-wide loss of DNA methylation (see Methods) (**Table S3**). Although this technique is less sensitive, it also shows a concentration dependent decrease in both AZA and ZEB up until 100 μM.

**Figure 1.**
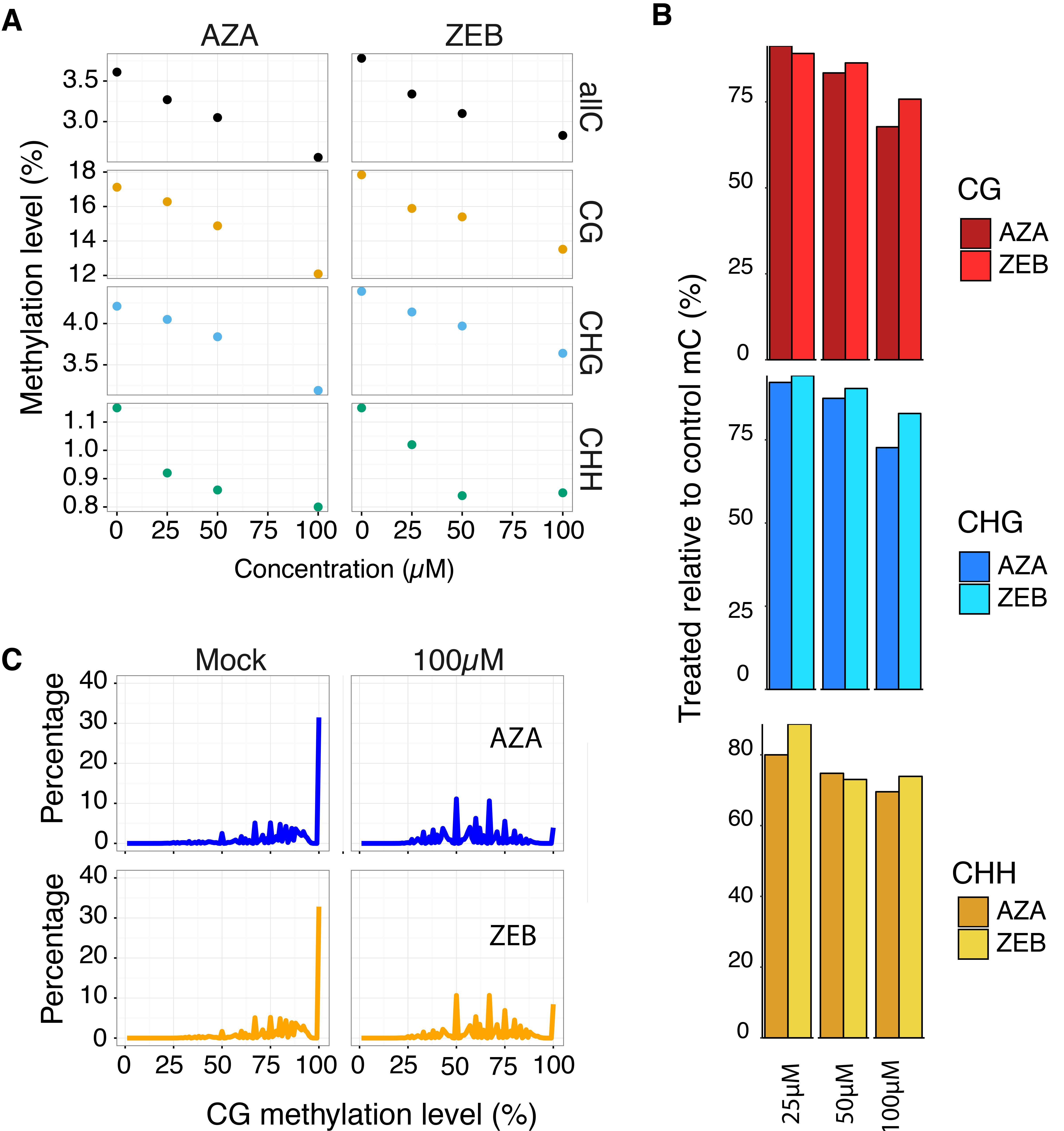
**AZA and ZEB treatment result in non-selective, concentration dependent loss of DNA methylation genome-wide**. A. The genome-wide methylation level of the control seedlings (0 μM) and seedlings treated with 25 μM, 50 μM, and 100 μM of either AZA of ZEB. B. The methylation level of AZA and ZEB-treated seedlings relative to the untreated control (treated/control) shown side-by-side for each context of DNA methylation. Both AZA and ZEB were compared to the untreated control methylation levels. C. A frequency distribution of the methylation levels of individual methylated cytosines for both AZA and ZEB treatments and the controls. In mock-treated samples, many methylated CG sites are 100% methylated, whereas in treated samples, most of the sites are not completely methylated.

To compare the genome-wide demethylating potential of each chemical, the methylation level of the treated samples relative to the control was plotted (**Figure 1B**). For consistency, each sample was compared to the untreated control (no DMSO added). The effect of the chemicals is similar but not identical, with either AZA or ZEB having a slightly greater effect at lower concentrations (25 μM and 50 μM). At a concentration of 100 μM, however, AZA had an 8.0% and 10.2% larger demethylating effect on mCG and mCHG, respectively. This was unexpected since the seedlings were treated for 10 days without replenishing the chemicals, and AZA has a far shorter degradation half-life than ZEB at room temperature (Champion et al. 2010). Both chemicals were found to reduce the distribution of methylation levels at highly methylated CG sites (**Figure 1C**), shifting the percentage of methylated CG dinucleotides that are completely methylated from 32.8% in the untreated control to 3.9% in 100 μM AZA-treated seedlings and 8.5% in 100 μM ZEB-treated seedlings.

To examine how AZA and ZEB affect methylation across chromosomes, the methylation level was calculated for 50 kb bins across chromosome 1 (**Figure 2A**). DNA methylation was reduced across chromosome 1 in a concentration-dependent manner, most notably for CG and CHG in the pericentromeric region. To illustrate the magnitude of demethylation along the genome, the methylation level of AZA and ZEB-treated DNA relative to that in the control DNA was plotted for each window across chromosome 1 (**Figure 2B**) and all chromosomes (**Figure S1**). CG methylation, maintained at higher levels than mCHG and mCHH across the genome (Zhang et al. 2006; Zilberman et al. 2007; Cokus et al. 2008; Lister et al. 2008), is consistently affected across the entire chromosome, whereas the loss of CHG and CHH methylation is greatest in the pericentromere. To eliminate bias from unmethylated regions and compare the demethylating effect in the pericentromere and chromosome arms, the genome was further broken up into 100 bp windows, and the 100 μM AZA-treated and mock-treated control methylation levels were plotted pairwise for both regions on chromosome 1 (**Figure 2C–Figure 2D**). Viewed this way, it becomes clear that AZA treatment affects highly methylated regions in both the pericentromere and chromosome arms equally, resulting in similar levels of demethylation. Given the evidence that RNA-independent DNA methyltransferases are the primary mediators of DNA methylation in the pericentromere, whereas gene body DNA methylation and RdDM activity are primarily found in the chromosome arms (Stroud et al. 2013; Zemach et al. 2013), this result suggests that chemical demethylating agents act without bias on the different pathways. Of note, windows with high CHH methylation (>25% methylation levels) tended to lose 50% to 75% of it in the AZA-treated sample (**Figure 2C**), whereas, CG methylation was less impacted, hovering between 25% and 50% loss (**Figure 2D**). Similar results were found in analyzing the 100μM ZEB-treated sample against the untreated control (**Figure S3**).

**Figure 2.**
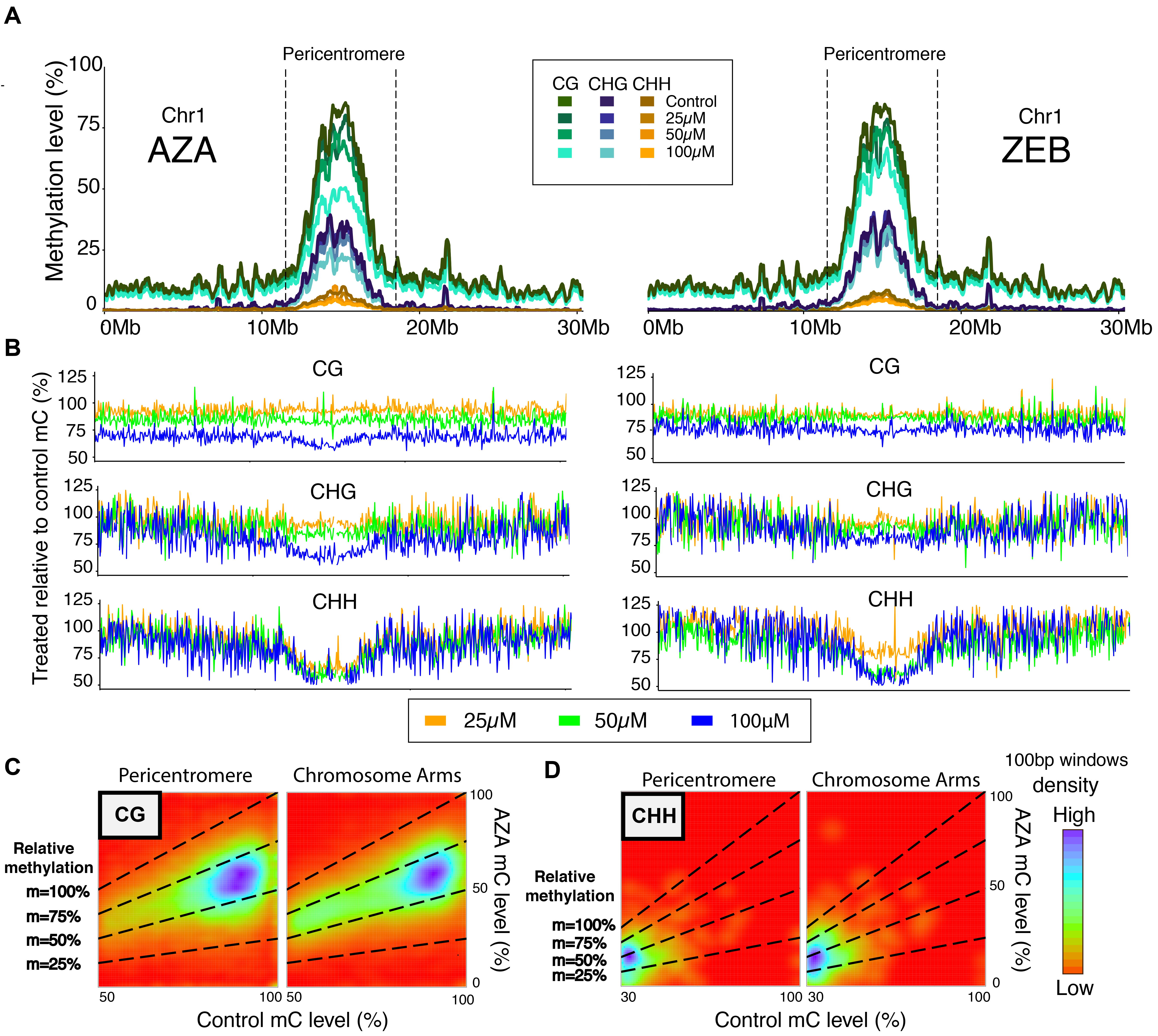
**AZA and ZEB cause a concentration-dependent loss of DNA methylation across chromosome 1**. A. The methylation level (all contexts) for each discrete 50 Kb window across chromosome 1 shown for untreated control samples and each treatment concentration of both AZA and ZEB. The dashed lines partition 7.5 Mb of area that was defined as the pericentromeric region of the chromosome. Refer to the legend for the concentration and context of methylation that each line represents. B. The methylation level of AZA-treated (left) and ZEB-treated (right) DNA relative to the mock-treated control is mapped across chromosome 1 for each 50 Kb region. C-D. A pairwise comparison of methylation level in mock-treated control seedlings and 100 μM AZA-treated seedlings for highly methylated 100 bp windows in both the pericentromere and the chromosome arms (as defined in 3A). CG (C) and CHH (D) contexts of DNA methylation are shown. A highly methylated window was defined as having >50% methylation in the control sample for CG and ≥30% methylation in the control sample for CHH. AZA-treated seedling methylation level is on the y-axis and control methylation level is on the x-axis. The color spectrum—ranging from red (low) to purple (high)— illustrates the density of points at a coordinate. The slopes (m) of the dashed lines represent the following relative methylation levels: 100% (treated and control methylation level are the same), 75%, 50% (treated methylation level is half of the control), and 25%.

The associated effect of DNA methylation on genes depends upon the methylation profile within the genes. Genes that are only methylated in the CG context are known as gene body methylated (gbM) and these genes are often expressed and moderate levels (Tran et al. 2005; Zhang et al. 2006; Zilberman et al. 2007). This is in contrast to genes that are methylated in all contexts as they are associated with lower gene expression or silencing (Law and Jacobsen 2010). Genes were previously classified into one of four classes based on their methylation profile (Niederhuth et al. 2016). Coding sequences of gbM genes are methylated in the CG context only. CHG-enriched (mCHG) genes contain significant numbers of methylated CHG, but not CHH sites, whereas CHH enriched (mCHH) genes contain significant numbers of methylated CHH sites and are typically methylated in all sequence contexts. To investigate the effect of AZA and ZEB on these different gene classes, average methylation was plotted across the gene bodies and 1500 base pairs up and downstream. In gbM genes, DNA methylation at CGs is reduced in a concentration-dependent manner for both AZA and ZEB (**Figure S2A**). Similarly, mCHH genes and transposons show a concentration-dependent loss of DNA methylation in all contexts (**Figure S2C-Figure S2D**). Comparing the 100 μM AZA and ZEB samples reveals that DNA methylation in all sequence features is more reduced by AZA, except for the CHH context in mCHH genes (**Figure 3A–Figure 3C**). Similarly, the difference in methylation level of AZA and ZEB-treated seedlings across transposons (**Figure 3C**) is less drastic for CHH methylation. This could suggest that at 100 μM, the effect of each drug is saturated.

**Figure 3.**
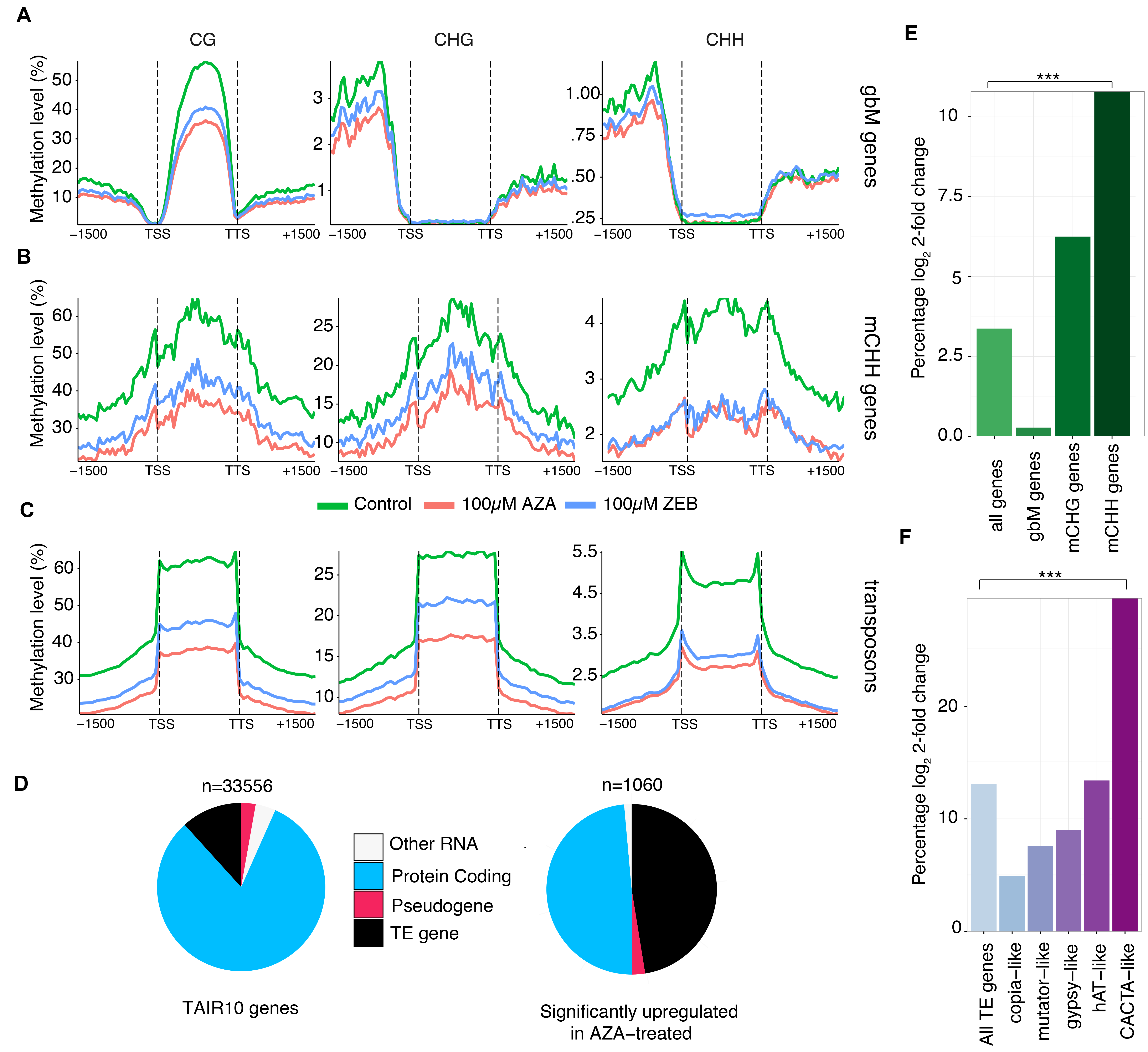
**AZA and ZEB cause similar patterns of DNA methylation loss and increase expression of genes highly methylated in all contexts**. A-C. The methylation level across all gene-body, CHH-enriched (mCHH), and TE genes are depicted for 100 μM AZA (red), 100 μM ZEB (blue), and untreated control (green). A pie chart depicting the types of genes upregulated by AZA treatment (right) compared with all genes annotated by TAIR10 (left). E-F. The percent of genes (protein-coding and TE) that are significantly upregulated when treated with 100 μM of AZA is shown. Statistical enrichment based on a Fischer two-sided test is denoted by ^*^ for *p* value < 0.05, ^**^ <0.01, and ^***^ < 0.001.

Methylation in all three contexts is often indicative of RdDM (Law and Jacobsen 2010) and can reduce gene expression of reporter genes (Hohn et al. 1996; Dieguez et al. 1997). Furthermore, previous studies have demonstrated that both AZA and ZEB reactivate transcription of silenced genetic elements in plants (Chang and Pikaard 2005; Baubec et al. 2014). To investigate the impact of DNA methylation inhibitors on the different methylation-based gene categories, we performed RNA-seq on seedlings treated with 100 μM of AZA. Out of all genes, 1060 were significantly upregulated and 263 were significantly downregulated in comparison to the mock-treated control (**Table S4**). Of these, 516 were protein-coding genes and 503 were classified as transposable element (TE) genes, a disproportionate amount when compared to the totality of genes annotated in the *A. thaliana* genome (**Figure 3D**). Of protein-coding genes, mCHH genes were found to be significantly enriched in upregulated genes based on a Fischer’s Exact Test (*p* value < 2.2 × 10^−16^), whereas mCHG and gbM genes were not significantly enriched (Figure 3E, **Table S4**). To further investigate effects on TE genes, differential expression of individual TE families was examined (**Figure 3F**). Out of all TE genes, 12.9% were upregulated. Copia-like elements were the least affected, with only 23 out of 491 genes (4.68%) upregulated. In contrast, the most highly upregulated category of TE genes was CACTA-like elements (29.8%) and they were the only category of TE genes that was significantly enriched when compared to all other TE genes (*p* value = 7.04 × 10^−12^, Fisher’s Exact Test).

Having investigated methylation-based gene classes and TE genes, we next examined specific genes, *FLOWERING WAGENINGEN (FWA)* and *SUPPRESSOR of drm1 drm2 and cmt3 (SDC)*, that do not fit into our methylation-based categories and nonetheless are known to be transcriptionally silenced by DNA methylation (Soppe et al. 2000; Kinoshita et al. 2007; Henderson and Jacobsen 2008). Although methylation was not completely lost, methylation levels were reduced, without bias to the sequence context (**Figure 4A–Figure 4D**). This is similar to what was observed genome-wide. Whether this is the result of a complete loss of DNA methylation in some cell types, which leads to reactivation of transcription, or if there is partial loss of DNA methylation in all cell types, remains to be investigated. *FWA* and *SDC* were among the top 10% of AZA-upregulated genes, with mRNA expression increased 5.4-fold and 6.9-fold respectively (**Table S5**). These well-studied genes, where the association of gene expression and epigenetic silencing has been established, along with the increased expression of many mCHH genes, provides further evidence that perturbation of DNA methylation by inhibitors predictably reactivates certain genetic elements.

**Figure 4.**
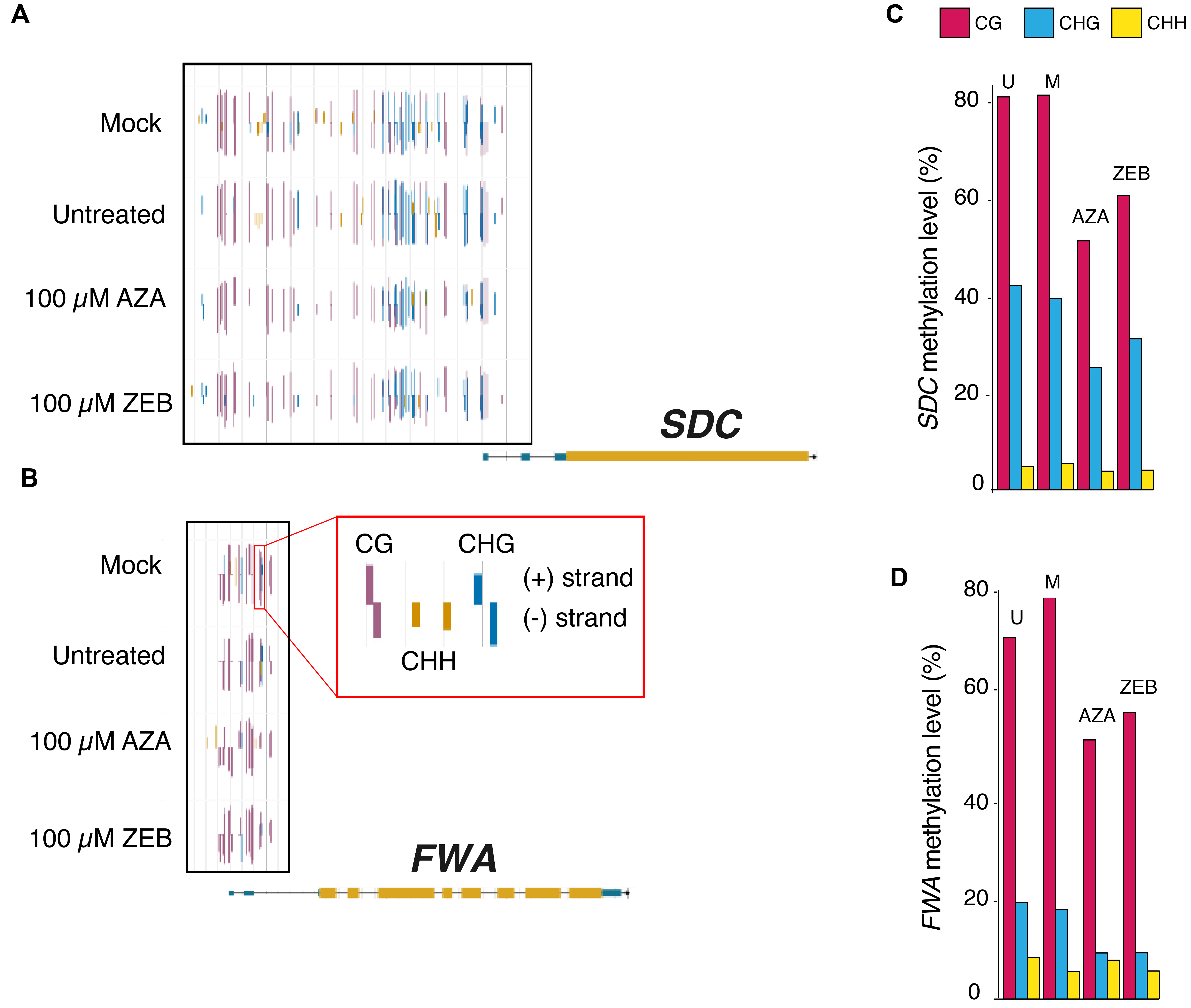
**FWA and SDC methylation level decreases and mRNA expression increases in response to chemical demethylating agents**. A-B. Browser screenshots of the methylation mapped to the *A. thaliana* genome show single base pair resolution data on individual cytosines for *SDC* and FWA. The legend box (outlined in red) shows that the height and the color of the bar illustrate the methylation level and context of each cytosine, whereas the direction of the bar indicates the strand. The genes themselves are mapped below the methylation data with the UTRs in blue, coding regions in yellow, and introns as the black line connecting them. C-D. The total methylation level (left) of the 5’UTR and upstream promoter region of genes *SDC* and *FWA* are depicted (U=Untreated, M=Mock). The black box in the browser screenshots surrounds the region analyzed for each gene.

## Discussion

In this study, we have examined the genome-wide effects of the chemical demethylating agents 5-azacytidine and zebularine. Known to have analogous modes of action (Pecinka and Liu 2014), our analysis demonstrates that both chemicals have similar effects, as DNA methylation is depleted across the entire genome in all sequence contexts. In previous studies, it had been estimated that at 40 μM, ZEB is a more effective demethylating agent than AZA (Baubec et al. 2009). Our estimates of relative methylation loss show that AZA may have a larger effect than ZEB at higher concentrations, whereas at lower concentrations, the difference is less clear. The differences in these results between labs may be explained by differences in growth medium composition, treatments of the plant material, growth conditions and duration of treatment. It could also suggest that certain loci are more susceptible at lower concentrations than others with regards to transcriptional reactivation. In addition, we found that highly methylated areas of pericentromeres and chromosome arms are comparably affected by demethylating agents and that CHH methylation is proportionally more impacted than CG or CHG by AZA. This could be due to indirect effects, as CHH methyltransferases rely in part on CG and CHG methylation (Stroud et al. 2013). Alternatively, at high concentrations, AZA may have a greater initial effect that persists over cell division.

We find that when *A. thaliana* is treated with either AZA or ZEB, there is a comparable concentration-dependent loss of DNA methylation for all sequence contexts across gbM genes, mCHH genes, and transposons. RNA-seq data revealed that these chemicals have a disproportionate transcriptional impact on mCHH genes and transposons. The enrichment for mCHH, as opposed to mCHG, gene reactivation suggested that transcriptionally silenced genes contain high levels of all three contexts of methylation. Among TE genes, CACTA-like elements were the the most highly upregulated. This category of mobile element is primarily localized in the pericentromeric region of chromosomes (Miura et al. 2004) and they have been found to be upregulated in *ddm1* mutants (Jeddeloh et al. 1999; Brzeski and Jerzmanowski 2003). Their low copy number and chromosomal position in *A. thaliana* hint that their expression is likely suppressed due to the deleterious effects of their transposition (Miura et al. 2004). Further, although it has been shown that DNA methylation is largely recovered in adult plant tissue after treatment with methylation inhibitors (Baubec et al. 2009), the lasting effect of treatment may go beyond each plants epigenetic profile. After inhibitor treatment, if transposition occurs in the germline or somatic tissue cells that are germline-progenitors, then any TE-inflicted mutations would be passed to subsequent generations. WGBS experiments on the offspring of inhibitor-treated plants could help answer questions about increased TE gene transposition and the impact of new insertions on DNA methylation of surrounding DNA.

Although AZA has been shown to be less stable than ZEB (Pecinka and Liu 2014), it causes approximately the same magnitude of DNA methylation loss genome-wide after 10 days of seedling growth and appears to have a greater effect at high concentrations. This may be due to ZEB incorporating less frequently into the DNA double helix (Jones and Taylor 1980; Ben-Kasus et al. 2005; Liu et al. 2015) or binding less strongly to DNA methyltransferases (Champion et al. 2010). Indeed, a recent assay of ZEB-treated *A. thaliana* did not detect deoxyzebularine (the processed, deoxyribonucleotide version of ZEB) in DNA at a sensitivity of ~1 deoxyzebularine to 5000 deoxycytidine, showing that it does not incorporate into DNA efficiently (Liu et al. 2015). Furthermore, the primary DNA repair pathways that are activated in ZEB and AZA-treated plants were shown to differ. Nucleotide excision is the dominant pathway in the of repair AZA-induced DNA damage, while homologous recombination was found to mainly mediate ZEB-induced damage repair (Liu et al. 2015). Any difference in the rate at which these nucleotide analogs are removed from the DNA helix may contribute to a difference in the amount of inhibitor-caused demethylation.

Here we demonstrate that AZA and ZEB treatment of *A. thaliana* results in similar changes in DNA methylation across the genome. In some ways, this is unexpected, given the evidence they may incorporate into DNA differently and largely be repaired by different pathways. Although some difference in the magnitude of DNA methylation loss may exist between AZA or ZEB-treated plants, the biological impact of the disparity is not yet known. Having used aggregated WGBS data from a population of cells, there is no easy way of knowing the effect on specific individual cells. Furthermore, experimental comparison is needed to determine if such a difference is impactful or whether the difference is biologically negligible. This study provides a detailed look into the genome-wide and transcriptional effects of commonly used DNA methylation inhibitors.

## Acknowledgements

We would like to thank Ortrun Mittelsten Scheid for critical comments and suggestions on this study. This work was supported by the National Science Foundation (NSF) (MCB - 1402183), by the Office of the Vice President of Research at UGA, and by The Pew Charitable Trusts to R.J.S. C.E.N was supported by a NSF postdoctoral fellowship (IOS - 1402183).

## Figure Legends

**Figure S1 | DNA methylation across all chromosomes is decreased when treated with AZA and ZEB in a concentration-dependent fashion**.

A-B. Relative methylation level shown across chromosomes 2-4 (chromosome 1 shown in Figure 2B) for AZA-treated (S1A) and ZEB-treated (S1B) seedlings.

**Figure S2 | AZA and ZEB induce a concentration-dependent decrease in DNA methylation in all types of genetic elements**.

A-D. DNA methylation maps across four different categories of methylated genetic elements: gbM genes (A), CHG-enriched (mCHG) genes (B), CHH enriched (mCHH) genes (C), transposons (D). The colors (top) represent the different treatment concentrations. The demethylating agent and sequence context is given above each vertical column of graphs.

**Figure S3 | Pairwise comparison of highly methylated 100 bp windows between ZEB-treated and control seedlings**.

A pairwise comparison of methylation level in untreated seedlings and the 100 μM ZEB-treated seedlings for highly methylated 100 bp windows in both the pericentromere and the chromosome arms (as defined in Figure 2A). CG and CHH contexts of DNA methylation are shown. A highly methylated window was defined as having ≥50% methylation in the control sample for CG and ≥30% methylation in the control sample for CHH. ZEB-treated seedling methylation level is on the y-axis and control methylation level is on the x-axis. The color spectrum — ranging from red (low) to purple (high)— illustrates the density of points at a coordinate. The slopes (m) of the dashed lines represent the following relative methylation levels: 100% (treated and control methylation level are the same), 75%, 50% (treated methylation level is half of the control), and 25%.

